# Histology-guided mathematical model of tumor oxygenation: sensitivity analysis of physical and computational parameters

**DOI:** 10.1101/2024.03.05.583363

**Authors:** Awino Maureiq E. Ojwang’, Sarah Bazargan, Joseph O. Johnson, Shari Pilon-Thomas, Katarzyna A. Rejniak

## Abstract

A hybrid off-lattice agent-based model has been developed to reconstruct the tumor tissue oxygenation landscape based on histology images and simulated interactions between vasculature and cells with microenvironment metabolites. Here, we performed a robustness sensitivity analysis of that model’s physical and computational parameters. We found that changes in the domain boundary conditions, the initial conditions, and the Michaelis constant are negligible and, thus, do not affect the model outputs. The model is also not sensitive to small perturbations of the vascular influx or the maximum consumption rate of oxygen. However, the model is sensitive to large perturbations of these parameters and changes in the tissue boundary condition, emphasizing an imperative aim to measure these parameters experimentally.

## 1. INTRODUCTION

Tortuous tumor vasculature can cause heterogeneities in tissue oxygenation, resulting in well-oxygenated areas (normoxia) and regions with low levels of oxygen (hypoxia) within a tumor [1]. Experimental measurements, including vascular parameters such as perfusion and hemoglobin-oxygen saturation, direct oxygen measurements with probes and oximetry, and markers such as immunohistochemistry [2], provide insight into the oxygenation status of tumor vs. normal tissues. However, a noninvasive and inexpensive approach to predicting tissue oxygenation is to recreate it using the first principles of physics and simulations of mathematical models of oxygen kinetics defined on a cellular microscale level [3]. This approach takes advantage of the spatial distributions of vessels and cells, which are assessed from tissue histology images, and treats them as agents in a hybrid agent-based model (ABM) [4, 5]. This enables the modeling of individual cell–cell interactions and cellular heterogeneity, such as differences in cell size, shape, and spatial location. Hybrid models combine discrete equations that describe the behavior of these agents with continuous equations that describe the kinetics of microenvironmental factors, such as diffusible oxygen and nutrients [4].

Input parameters in mathematical models, including the hybrid agent–based models, may be associated with some uncertainty that can produce a range of possible model outputs. Sensitivity analyses can be used to understand the effect of an input parameter’s perturbation on the model output and to identify the robust parameters whose perturbation does not change the model outputs significantly [6]. Sensitivity analysis also identifies the parameters contributing to prediction inaccuracy in the model output; that is, it shows whether changes in the input parameters’ values significantly alter the model output [6]. It is vital to assess how sensitive the model outputs are to small changes to the input parameter values, as this can yield insight into the model output changes attributed to the dynamics of the modeled biological system vs. those attributed to artifacts from the unknown aspects of the biological system (biological uncertainty) or parametrization [7, 8]. Sensitivity analysis involves local and global techniques. For local analysis, the input parameter values are changed one at a time while the other parameters remain fixed. For global analysis, all model parameters are changed simultaneously.

There are several methods for sensitivity analysis of agent-based models, as discussed in [9]. Here, we focus on robustness sensitivity analysis, which is a type of local sensitivity analysis investigating the robustness of a model’s output to local perturbations of input parameters. We apply this analysis to parameters related to the initial oxygen level in the tissue, the computational and tissue boundary conditions, the influx of oxygen from vessels, and cellular uptake. Our analysis provides insight into which model parameters can be varied and also identifies the parameters whose accurate values would need to be measured experimentally.

The rest of the paper is organized as follows. The mathematical model is described in **section 2**. The results presented in **section 3** include the design of the digitized tissue **(section 3.1)**, the calculation of a stable oxygen distribution **(section 3.2)**, a sensitivity analysis of the model initial conditions **(section 3.3)**, the domain boundary conditions **(section 3.4)**, the tissue boundary conditions (**section 3.5)**, vascular influx **(section 3.6)**, and the Michaelis–Menten parameters for cellular uptake **(section 3.7)**. Finally, we discuss the implications of our sensitivity analyses **(section 4)**.

## 2. MATHEMATICAL MODEL

We used a hybrid multicell lattice-free agent-based model that includes individual cells and vasculature represented as discrete agents, with a continuous description of oxygen kinetics, similar to our previous models [3, 10, 11]. However, here we used the digitized tissue obtained from a histology image as our modeling domain (section 2.1), which requires preprocessing steps to resolve cell–cell and cell–vessel overlaps (section 2.2), to annotate the outer points (section 2.3), and to annotate the inner cavities (section 2.4) for tissue boundary conditions. The influx of oxygen from individual vessels was also modeled as a boundary condition (section 2.5), and a reaction–diffusion partial differential equation was used to describe the oxygen kinetics (section 2.6). The cellular uptake by tumor and immune cells was governed by Michaelis–Menten kinetics (section 2.7). Finally, the sensitivity analysis methods we used are described in section 2.8.

### 2.1. Data acquisition

Our mathematical model uses tissue histology images as a computational domain. These images were acquired from *in vivo* experiments with C57BL/6 female mice that received an intravesical instillation of 1×10^5^ MB49-OVA murine bladder cancer cells. At the end of the experiment, the bladder tissue was harvested and sliced into sections 4 μm thick and mounted on slides. The sections were stained with hematoxylin and eosin (H&E) to visualize all cells’ nuclei and cytoplasm, and immunohistochemistry (IHC) stains specific for vasculature (CD31). The slides were then scanned with a Leica Aperio AT2 digital pathology slide scanner, and the H&E and IHC images were co-registered using Visiopharm’s Tissuealign. Visiopharm was used to segment the H&E image images. The cell detection algorithm was used to segment the cells and vessels and determine their coordinates and areas. The segmented H&E image was used to identify domain and tissue boundaries using the minimum and maximum coordinates of all cells. The shapes of cells and vessels were approximated by circles. Thus, we calculated the circle radius of each cell and vessel using Eq. (1).

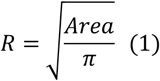

### 2.2 Resolve cell–cell and cell–vessel overlaps

Using circles to approximate the cell and vessel shapes may cause overlaps between them. Therefore, the repulsive forces were applied between nearby cells (cell–cell forces) and between cells and vessels (cell– vessel forces) to resolve overlaps. Let ***X***_*i*_ and ***X***_*j*_ be the coordinates of two discrete elements (either cells or vessels) with radii *R*_*i*_ and *R*_*j*_, respectively. The Hookean force 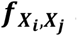 acting on element ***X***_*i*_ is given by Eq. (2),

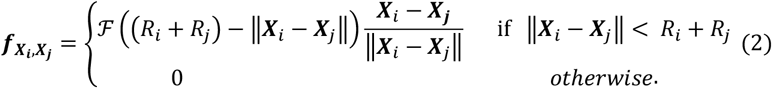

where *ℱ* is a constant spring stiffness identical for all cells and vessels and *R*_*i*_ + *R*_*j*_ represents the force resting length.

The repulsive force 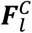 acting on cell 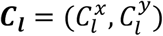 combines force contributions from all neighboring cells 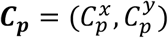 and vessels 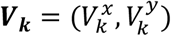 and is given by Eq. (3), where *N*_*c*_ represents the number of cells and *N*_*v*_ represents the number of vessels.

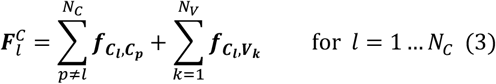

Cell relocation to resolve cell–cell and cell–vessel overlaps follows the overdamped spring equation:

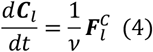

where ν is the viscosity of the surrounding medium. We applied Eqs. (3), and (4) iteratively to all overlapping cells until the magnitude of the repulsive force 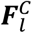 fell below a small threshold value *F*^*max*^, at which point the remaining cell–cell and cell–vessel overlaps are negligible. However, vessel–vessel overlaps are allowed as they mimic the different vascular shapes observed in histology images that arise when vessels within the tissue slice are cut at different angles. This algorithm to resolve overlaps was used once during the preprocessing step in order to create the digitized tissue for further oxygen simulations. The force algorithm was deactivated after this step.

### 2.3. Define outer points

As our computational domain is rectangular and tissue histology can have an irregular shape that does not conform to that domain, we identified a tissue boundary to separate grid points outside and inside the tissue. The tissue boundary was defined using MATLAB® *boundary* routine based on the cell coordinates after resolving overlaps. We determined the grid points located outside the tissue boundary called “outer points” using the MATLAB® *inpolygon* function. This function takes the tissue boundary coordinates and the grid points as input values and returns the value one if the boundary encloses the given grid point and zero if it does not. The outer points were used to define tissue boundary conditions to avoid the calculation of oxygen diffusion outside the tissue.

### 2.4. Define inner cavities space

The bladder is a hollow organ lined by a luminal mucosa. Thus, our computational domain included grid points that represented empty inner cavities. To avoid calculating the diffusion of oxygen into these empty spaces, we needed to specify the boundary conditions at all cavity points. However, the bladder tissue (especially normal, non-tumor tissue) can be composed of sparsely packed cells that may be misclassified as cavity points. To distinguish between these two types of points, we introduced “ghost cells” to artificially fill the spaces around cells segmented from a histology image. The ghost cells remove the sparsely packed cells; thus, the remaining points with no cells (real or ghost) in their vicinity belong to inner cavities.

Ghost cells were created following Eq. (5). For every sparsely packed cell ***C***_***l***_, that is, a cell for which the minimum Euclidean distance from other cells was greater than a specified threshold *Th*_*G*_, ten ghost cells were placed in a random direction *θ*_*i*_ (for *i*=1…10).

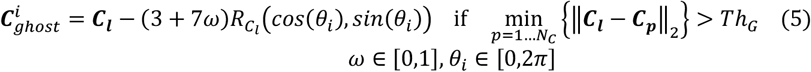

Similarly, the ghost cells were added around each vessel ***V***_***k***_ using Eq. (6) if the minimum distance from its closest cell was greater than *Th*_*G*_.

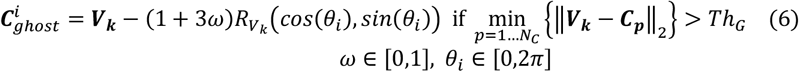

Thus, a grid point was defined as belonging to an inner cavity only if its minimum Euclidean distance from all cells, ghost cells, and vessels was greater than a specified threshold *Th*_*C*_. This process was repeated for all grid points inside the tissue boundary to identify those that form all inner cavities. Identification of all inner cavity points was done as a one-time preprocessing step to reduce computational costs.

### 2.5. Define vascular influx

Since our goal was to recreate tissue oxygenation based on a static histology image, we assumed that the vessels would not evolve during the simulation and that the oxygen level in each vessel was also not changing. Therefore, we could identify the grid points inside each vessel and specify the oxygen influx values at each point. This one-time preprocessing step allowed us to use these values in every consecutive iteration. The computational domain was divided using the grid size *h* equals 5 μ*m*. Because the vessels identified from the histology image may have various diameters, we considered two cases: the vessels with diameters (i) larger than the grid size and (ii) smaller than the grid size. For large vessels (such that, 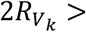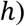, we assumed that there was a constant amount of oxygen (*I*_*γ*_) supplied at each grid point (*x*_*i*_, *y*_*j*_) inside the vessel with center at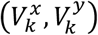; thus, the level of oxygen at the grid point γ_*i,j*_ = γ(*x*_*i*_, *y*_*j*_) and the total amount of oxygen in the vessel γ(***V***_***k***_, *t*) are given by Eq. (7)

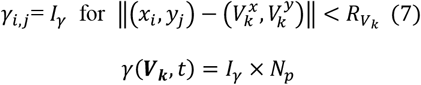

where *I*_*γ*_ is the defined oxygen level and *N*_*p*_ is the number of grid points inside the vessel ***V***_***k***_.

For the small vessels with diameters below the grid size and no grid points located inside the vessel, the oxygen level supplied from that vessel (*I*_*γ*_) needed to be distributed to the four grid points surrounding the vessel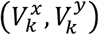. This was done so as to ensure that the oxygen level at each of the grid points was inversely proportional to its distance from the vessel center, as described in Eq. (8)

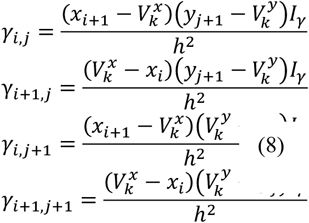

where γ_*i,j*_ + *γ*_*i*+1,*j*_ + γ_*i,j*+1_ + γ_*i*+1,*j*+1_ = *I*_*γ*_ and *I*_*γ*_ is the total vascular level of oxygen. The four grid points surrounding the vessel are (*x*_*i*_, *y*_*j*_), (*x*_*i*+1_, *y*_*j*_), (*x*_*i*_, *y*_*j*+1_), and (*x*_*i*+1_, *y*_*j*+1_).

In principle, it would be possible to reduce the grid size below the smallest cell radius, such that the grid points could be located inside every vessel. However, this was computationally too expensive. The grid points inside the large vessels and surrounding the small vessels were identified once as a preprocessing step, and the assigned appropriate influx values were used in all subsequent iterations of our algorithm. This was a one-time preprocessing step to reduce the computational cost.

### 2.6 Simulate tissue oxygenation

To recreate the tissue oxygenation landscape based on a static histology image, we generated a stable distribution of oxygen that was a result of the balance between oxygen influx, its diffusion, and cellular uptake. To achieve this, we ran the reaction–diffusion equation iteratively until the difference between consecutive oxygen distributions was very small.

The change in oxygen level at location ***x*** = (*x, y*) at time *t* depends on the influx from vessels (modeled as a boundary condition), diffusion through the tissue with a constant diffusion coefficient *D* (solved using the standard finite difference methods), and uptake by cells (governed by Michaelis–Menten kinetics). The overall oxygen kinetics is described in Eq. (9)

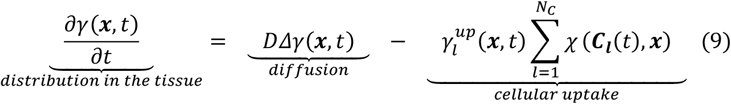

where 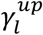 is the cellular uptake rate defined separately for small and large cells (see Eqs. (13) and (15)),

*N*_*C*_ is the total number of cells, and ***C***_*l*_ represents the cell coordinates. The oxygen is defined on the cartesian grid ***x*** = (*x, y*), and the cells are defined on the Lagrangian grid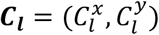. Therefore, the oxygen– cell interactions are specified by the indicator function, χ (***C***_***l***_(*t*), ***x***) given in Eq. (10) with the interaction radius 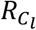 for large cells and interaction width *h* for small cells.

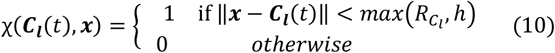

The initial oxygen level in the whole tissue area was set to zero. The following boundary conditions were set up:

i. in all grid points inside inner cavities and outside the tissue boundary, the oxygen level was set up to a hypoxia threshold γ_*h*_
ii. in all grid points representing the vessels, a constant level of oxygen was imposed according to Eq. (7) and Eq. (8)
iii. no loss or gain of oxygen along the domain boundaries was imposed, as defined by the Neumann-type boundary conditions:

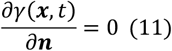

where ***n*** is an outward normal vector.

To achieve a stable distribution of oxygen within the tissue, that is, a balance between oxygen influx, diffusion, and cellular uptake, the reaction-diffusion equation was iteratively repeated until the L_2_norm between two consecutively calculated oxygen distributions fell below the prescribed threshold, i.e., 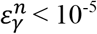, where:

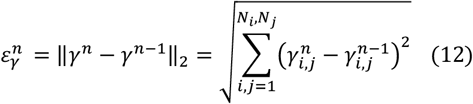

Once this criterium was met, the oxygen distribution within the tissue was considered numerically stable.

### 2.7. Define cellular uptake

Similar to the vessels described before in section 2.5, cells in the tissue may have diameters larger or smaller than the grid width. For a large cell with the center at 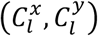 and diameter greater than *h*, there are grid points (*x*_*i*_, *y*_*j*_) inside the cells where the uptake of oxygen 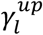 is defined by the Michaelis–Menten kinetics using Eq. (13) as,

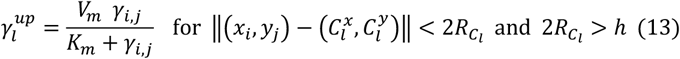

where γ_*i,j*_ is the oxygen level at grid point (*x*_*i*_, *y*_*j*_), 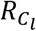 is the cell radius, *V*_*m*_ is the maximum consumption rate, and *K*_*m*_ is the oxygen level at half *V*_*m*_. The total cell oxygenation level γ(***C***_***l***_, *t*) in this case is defined by Eq. (14).

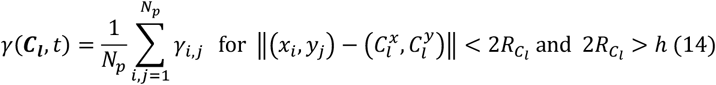

where *N*_*p*_ is the number of grid points inside the cell.

For a small cell 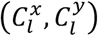 with no grid points inside and a diameter smaller than *h*, the cell will utilize the four surrounding grid points to absorb oxygen in amounts that are inversely proportional to the distance from the cell center. This is defined in Eq. (15).

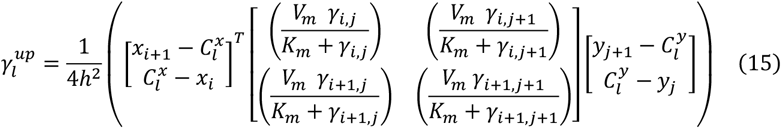

where γ_*i,j*_, γ_*i,j*+1_, γ_*i*+1,*j*_, *a*nd γ_*i*+1,*j*+1_, are the oxygen levels at the grid points (*x*_*i*_, *y*_*j*_), (*x*_*i*_, *y*_*j*+1_), (*x*_*i*+1_, *y*_*j*_), *a*nd (*x*_*i*+1_, *y*_*j*+1_) surrounding the cell. The total cell oxygenation level in this case is defined by Eq. (16).

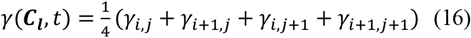

All parameter values are given in Table 1.

**Table 1.** The physical and computational model parameters.

| Parameters | Model value | Reference |
| --- | --- | --- |
| Force stiffness, $\mathcal{F}$ | 50 $\mu\text{g}/\mu\text{m.s}^2$ | [12, 13] |
| Medium viscosity, $\nu$ | 250 $\mu\text{g}/\mu\text{m.s}$ | [14] |
| Oxygen diffusion, $D$ | 100 $\mu\text{m}^2/\text{s}$ | [15, 16] |
| Baseline oxygen in a vessel, $I_\gamma$ | 65.28 $\sigma\text{g}/\mu\text{m}^3$<br>= 60 mmHg | [17, 18] |
| Hypoxia level, $\gamma_h$ | 10.8 $\sigma\text{g}/\mu\text{m}^3$<br>= 10 mmHg | [2] |
| Michaelis constant, $K_m$ | 1.344 $\sigma\text{g}/\mu\text{m}^3$<br>= $2.1 \times 10^{-3}$ mM | [19] |
| Maximum oxygen uptake rate, $V_m$ | 4.774 $\sigma\text{g}/\text{s}.\mu\text{m}^3$<br>= $3.25 \times 10^{-17}$ mole/cell/s | [19] |
| Repulsive force threshold, $F^{max}$ | 50 $\mu\text{g}/\mu\text{m.s}^2$ | |
| L<sub>2</sub> norm error threshold, $\varepsilon_\gamma$ | $10^{-5}$ | |
| Threshold for creating ghost cells, $Th_G$ | $1.5 \times h \mu\text{m}$ | |
| Threshold for creating inner cavities, $Th_C$ | $2.5 \times h \mu\text{m}$ | |
| Grid width, $h$ | 5 $\mu\text{m}$ | $\frac{D\Delta t}{h^2} \leq \frac{1}{4}$ |
| Time step, $\Delta t$ | 0.0625 s | |
| Scaling parameter, $\sigma$ | $0.5 \times 10^{-19}$ | |

### 2.8. Methods for sensitivity analysis

We followed [9] and considered here a sensitivity analysis method that determines how robust the model output is to local parameter perturbations, i.e., how changing the value of one parameter while holding the other parameter values fixed affects or does not affect the model output. The A-measure (section 2.8.1) was used as an intuitive way to compare the distributions produced by the perturbed parameter values vs. the baseline value [9]. L_2_ norm was used to determine the differences between two stabilized oxygen distributions (section 2.8.2).

#### 2.8.1. A-measure of stochastic superiority

The A-measure of stochastic superiority (or A-measure) is a method used to compare the equality of two data distributions (discrete or continuous) [20]. It describes the probability that a randomly chosen value from one distribution is greater than a randomly chosen member of the other distribution. It accounts for ties by awarding a value of 0.5.

Suppose we want to compare two distributions *B* = *b*_1_, *b*_2_, …, *b*_*m*_ and *C* = *c*_1_, *c*_2_, …, *c*_*n*_ with respect to some variable *X*. Using conventional probability theory, A-measure *A*_*BC*_ (*X*) is defined as,

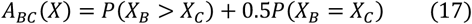

where *X*_*B*_ is a randomly selected value from *B* and *X*_*C*_ is a randomly selected value from *C*. The point estimate *Â*-measure is given by Eq. (18).

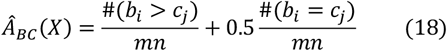

Hamis et al., 2020 [9] provided the following notation of *Â*measure:

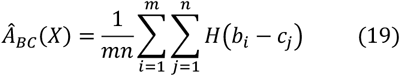

where *H*(*x*) is a Heaviside step function:

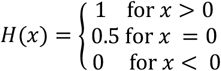

If *Â*_*BC*_ (*X*) = 0.5, the distributions *B* and *C* are stochastically equal with respect to *X*. Thus, *Â*-measure ∈ [0,1], assesses how much *B* and *C* deviate from the equality value, 0.5. Furthermore, the effect of the stochastic difference between *B* and *C* can be described using statistical significance based on the following classifications [20]: the statistical significance of the difference between *B* and *C* is small if *Â*-measure ∈ [0.44,0.56], medium if *Â*-measure ∈ [0.36,0.64], and large if *Â*-measure ∈ [0.29,0.71]. For example, if *Â*-measure is close to 0.5, the *B* and *C* distributions are “fairly equal”, and the statistical significance of the difference between *B* and *C* is classified as small. The *getA_measure*.*m* function used to implement A-measure in MATLAB® is provided in [9]. More details about A-measure’s derivation are described in [9, 20].

#### 2.8.2. L_2_ norm of differences between tissue oxygenation patterns

We defined the oxygen on a cartesian grid which we stored as matrices in MATLAB®. Calculating the L_2_ norms gives the difference between the two matrices containing oxygen values and quantifies whether the matrices are significantly different [3]. This study used L_2_ norms, calculated using Eq. (12), to determine the differences between the final stabilized oxygen distribution produced by the baseline parameter value vs. those produced by the perturbed parameter values. The calculations were done using only the tissue grid points and normalized by the number of tissue grid points. The lower the L_2_ norm value, the smaller the difference between the two distributions.

## 3. RESULTS

In this section, we describe how the digitized bladder tissue obtained from a scanned histology image (**section 3.1**) was used to recreate a stable oxygenation map using the hybrid ABM model (**section 3.2**).

The following local sensitivity analyses of model parameters were performed: sensitivity to initial conditions (**section 3.3**), to different computational domain boundary conditions (**section 3.4**), and to boundary conditions imposed on the inner cavities and outer points (**section 3.5**). The robustness of the vascular influx (**section 3.6**), the maximum cellular uptake *V*_*m*_ (**section 3.7**), and the Michaelis constant *K*_*m*_ (**section 3.8**) were also tested for small and large parameter perturbations. The number of perturbed parameter values was reported for each parameter investigated. Furthermore, one or more of the following model outputs were generated for each perturbed parameter value: 1) the average oxygen value in the tissue in each iteration, 2) the final stabilized oxygen distribution in the tissue, and 3) the cellular oxygen levels calculated using Eqs. (14) and (16) for each cell after stabilization of the oxygen gradient.

### 3.1. Histology data and digitized tissue

The H&E and CD31 histology images in Figs. 1A and 1B were used for cell and vessel segmentation, respectively. Next, the repulsive force algorithm was applied to resolve potential overlaps between cells and vessels. The final digitized tissue is presented in Fig. 1C. The tumor area is 0.75 mm^2^, containing 9,896.80 cells/mm^2^ and 745.66 vessels/mm^2^. The nontumor area is 11.88 mm^2^ with a cellular density of 3,682.66 cells/mm^2^ and a vascular density of 340.02 vessels/mm^2^. This results in the tumor area occupying 6% of the tissue and the nontumor area occupying 94%. These proportions, together with the sizes and spatial locations of cells and vessels, could influence the oxygen distribution in a tissue [3]. The tumor cells’ nuclei in this tissue were relatively small, with diameters of 5.191 ± 2.006 μ*m* (mean ± std). The nontumor cells nuclei (which may include immune and stromal cells) were smaller on average than the tumor cells (4.368 ± 1.461 μ*m*), while the vessels were relatively larger (13.737 ± 6.978 μ*m*). The tumor cells, nontumor cells, and vessels are shown in gold, gray, and red, respectively, in Fig 1C.

**Fig. 1.**
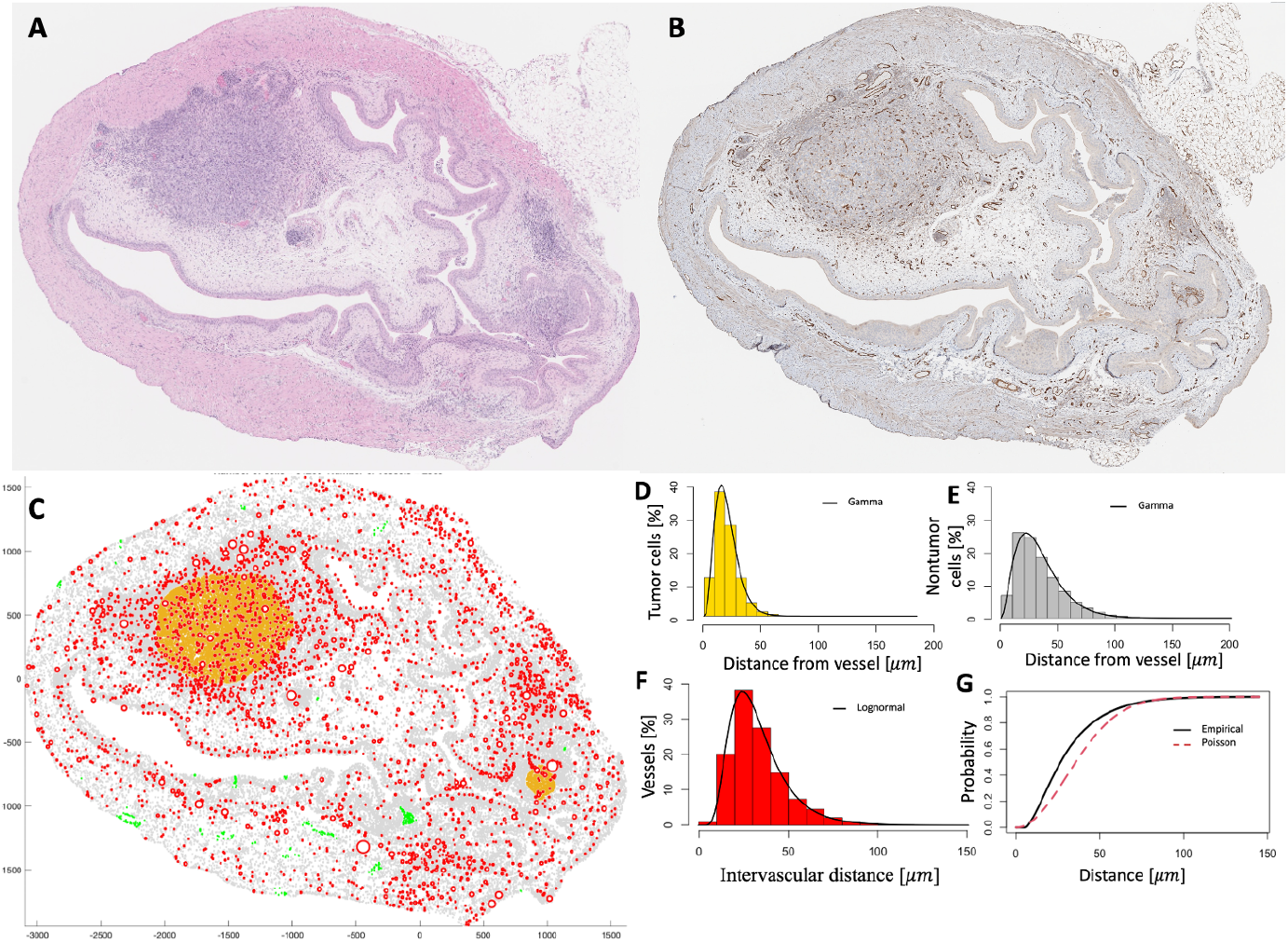
Histology data and the digitized tissue. **A**. A hematoxylin and eosin histology image of the tissue. The nuclei of tumor and nontumor cells are stained in deep blue-purple, and the different shades of pink represent the cytoplasm, extracellular matrix, and other structures. **B**. The CD31^+^ immunohistochemistry histology image with the endothelium of vessels stained in brown. **C**. The digitized tissue contains cells from A and vessels from B. The colors represent vessels (red), tumor cells (gold), and nontumor cells located within 100 μ*m* from the nearest vessel (gray), and nontumor cells that are more than 100 μ*m* away from a vessel (green). **D-E**. The frequency distributions of the minimum distances separating cells and vessels within the tumor and nontumor regions, respectively. **F**. The frequency distribution of intervascular distances in the tissue. **G**. A plot of the empirical G-function (black line) against the Poisson distribution (dashed red line) of the cells and vessels in the tissue.

Fig. 1D shows the relative frequency histogram of the minimum distances between cells and vessels in the tumor. This data is left-skewed and follows a gamma distribution fit (shape = 4.15, rate = 0.19), as indicated by the black line. This data indicates that 98% of the tumor cells are located within 50 μ*m* from the nearest vessel. Such cells can be expected to be well-oxygenated and were thus used for comparison to cell oxygenation in our simulations. Fig. 1E shows that most nontumor cells were also close to the vessels, except for 0.52 % of cells that were 100 μ*m* or more away from the nearest vessel. This minimum distance data also follows the gamma distribution fit (shape = 2.84, rate = 0.09), as indicated by the black line in Fig.1E. The cells that are more than 100 μm away from a vessel cells should be hypoxic [2] and were used to determine the baseline value of the maximum cellular uptake rate, *V*_*m*_ (section 3.7). The intervascular distances and spatial vascular distribution determine the available oxygen supply in a tissue [3]. Fig. 1F shows the relative frequency histogram of the minimum intervascular distances in the tissue. The data follows a lognormal distribution fit (mean = 3.4, std = 0.46, as indicated by the black line), showing that the most frequent separation was about 30 μ*m*. Of the vessels, 86% were less than or equal to 50 μ*m* from each other, and the remaining 14% were within 50 to 100 μ*m* of each other. As some vessels were closer to each other, regions with clustered vessels would be expected to be more oxygenated. We then performed a chi-square dispersion test of complete spatial randomness for the vessels based on quadrat counts using the *quadrat*.*test* in the R package *Spatstat*. This was done to determine whether the vessels were independently and homogeneously distributed in the whole tissue, meaning they did not differ in density (vessel number per area) from location to location. The function divides the domain into equal quadrats (a set of rectangles) and tests if the number of points in each quadrat deviates from a homogenous intensity [21]. The quadrat test indicated inhomogeneity of the vessels when 1x1 (p = 0), 10×10 (p = 0), or 100×100 (p = 2.78×10^−22^) array of equal quadrats were used.

Next, using the G-function, we tested whether the cells clustered around their nearby vessels in the tissue. The G-function calculates the nearest-neighbor probability, meaning the chance that a randomly chosen cell in the tissue has its nearest vessel within a given distance [22]. The G-function value can only increase up to a maximum of 1 because of the increasing radius around a given cell used in the calculations. The G-function values are compared to those of a completely random distribution (Poisson process) to determine the clustering or dispersion of the cells around the vessels. Fig. 1G gives the nearest neighbor probabilities (black solid line), as calculated by the multitype G-function, *Gcross*, in R package *Spatstat*.

The upward deviation from the Poisson line (red dashed line) shows that the nearest neighbor distances between cells and vessels are shorter at distances less than 75 μ*m* than would be expected if the pattern was completely random, indicating clustering. The cells and vessels were regularly spaced at distances greater than 75 μm.

### 3.2. Oxygen stabilization

The digitized tissue containing the locations of cells and vessels was used to recreate an oxygenation map of the bladder tissue. To achieve this, a balance was required between the continuous constant influx of oxygen from all vessels, oxygen diffusion through the tissue, and the continuous uptake by all tumor and nontumor cells. This balance was achieved through the stabilization process described by Eq. (12). Snapshots taken during the process of oxygen stabilization in the digitized tissue are shown in Fig. 2. The tissue was initialized with 0 mmHg of oxygen, which was uniformly distributed, as shown in Fig. 2A. Hypoxia at 10 mmHg of oxygen was imposed as a boundary condition in the outer points and inner cavities (Fig. 2A). Simulations were run iteratively, and the oxygen gradient was formed in the tissue as vessels continuously output oxygen at a constant rate; oxygen diffused through the tissue and was consumed by the cells via Michaelis–Menten kinetics. The oxygen maps in Figs. 2B and 2C show the difference in tissue oxygen gradients at iterations 100 and 4,500, respectively. As the number of iterations increased, the oxygen gradient eventually reached an equilibrium and stabilized (Fig. 2D), and subsequent changes in oxygen gradients were negligible. The simulations were stopped when the L_2_norm of the difference between subsequent oxygen distributions fell below the threshold value of 10^−5^. The final stabilized oxygen distribution reached an average level of 31.403 mmHg after 7,981 iterations (Fig. 2E), with ε_*γ*_= 9.94×10^−6^.

**Fig. 2.**
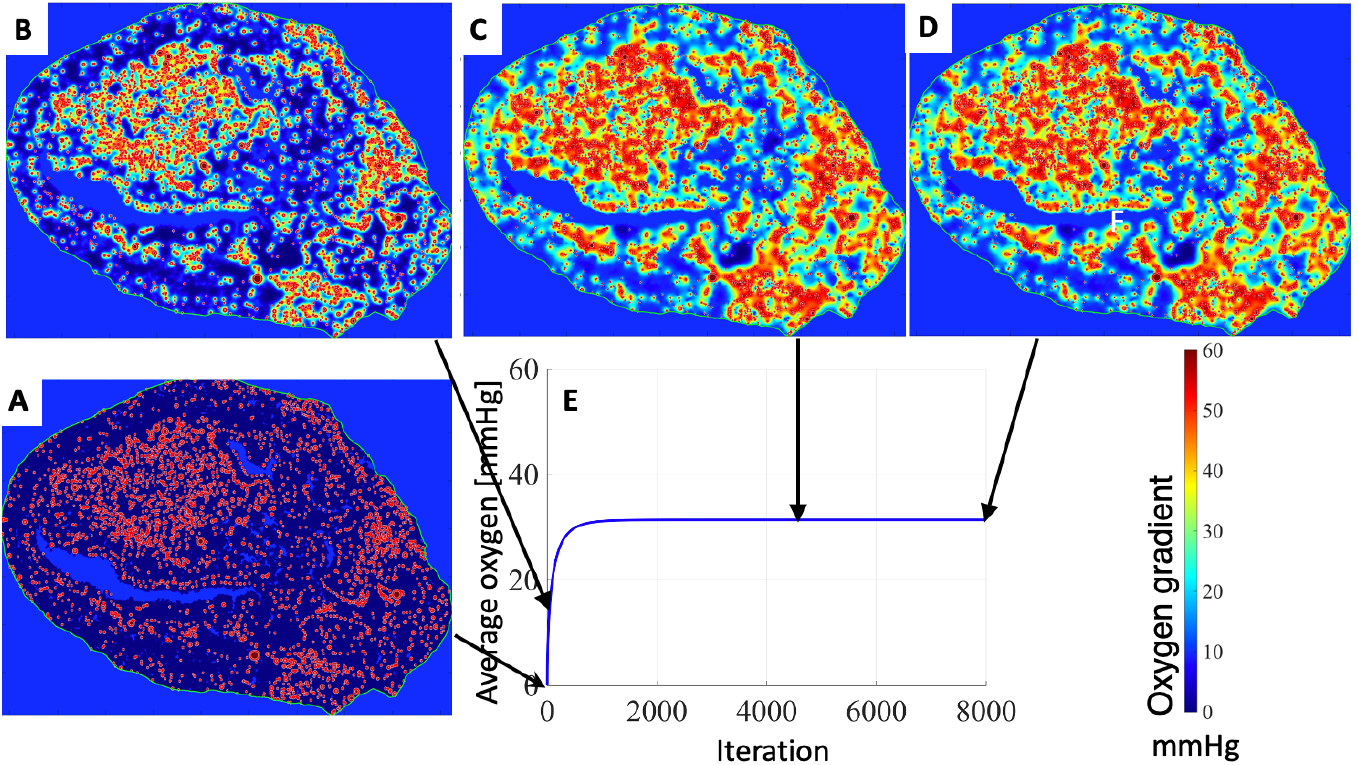
Tissue oxygenation during the stabilization process. **A**. The tissue initial setup with a uniform distribution of 0 mmHg of oxygen in the tissue and hypoxic values (10 mmHg) imposed in the inner cavities and the areas outside the tissue. The red circles represent the vessels. **B-C**. The simulated oxygenation maps after 100 and 4500 iterations, respectively. **D**. The numerically stable oxygenation map after 7981 iterations. **E**. Changes in the average oxygen level in the tissue (y-axis) over iterations (x-axis) from the initial 0 mmHg to the stable average level of 31.403 mmHg.

### 3.3. Sensitivity analysis of the initial tissue oxygenation

Initially, the oxygen was uniformly distributed within the tissue at a level of 0 mmHg. We tested whether varying the initial oxygen level from 0 mmHg to 60 mmHg with increments of 3 mmHg would impact the final stabilized oxygen distribution.

In cases when the initial level of oxygen in the tissue is low, oxygen must first outflux from the vessels and diffuse through the tissue before being consumed by the cells. Thus, the initial changes in the spatial distribution of oxygen were mostly due to the diffusion process. Fig. 3 shows that the average oxygen level rose before stabilizing for the simulations initialized with lower oxygen levels (blue curves). On the contrary, the average oxygen level decreased steadily until stabilization for simulations initialized with higher oxygen levels (red curves). This is because when the initial level of oxygen in the tissue was high (especially at 60 mmHg), the tissue was already oxygenated, and the cells could uptake the oxygen immediately; thus, the initial changes in the spatial distribution of oxygen were due mostly to the uptake process. Despite different initial oxygen levels, all 21 simulations stabilized at 31.403 mmHg (Fig. 3). Therefore, the model was insensitive to the level of oxygen chosen at the beginning of each simulation. However, the time required for the stabilization process to be completed differed for all cases. The simulations that started with lower oxygen levels stabilized faster than those with higher initial oxygen levels.

**Fig. 3.**
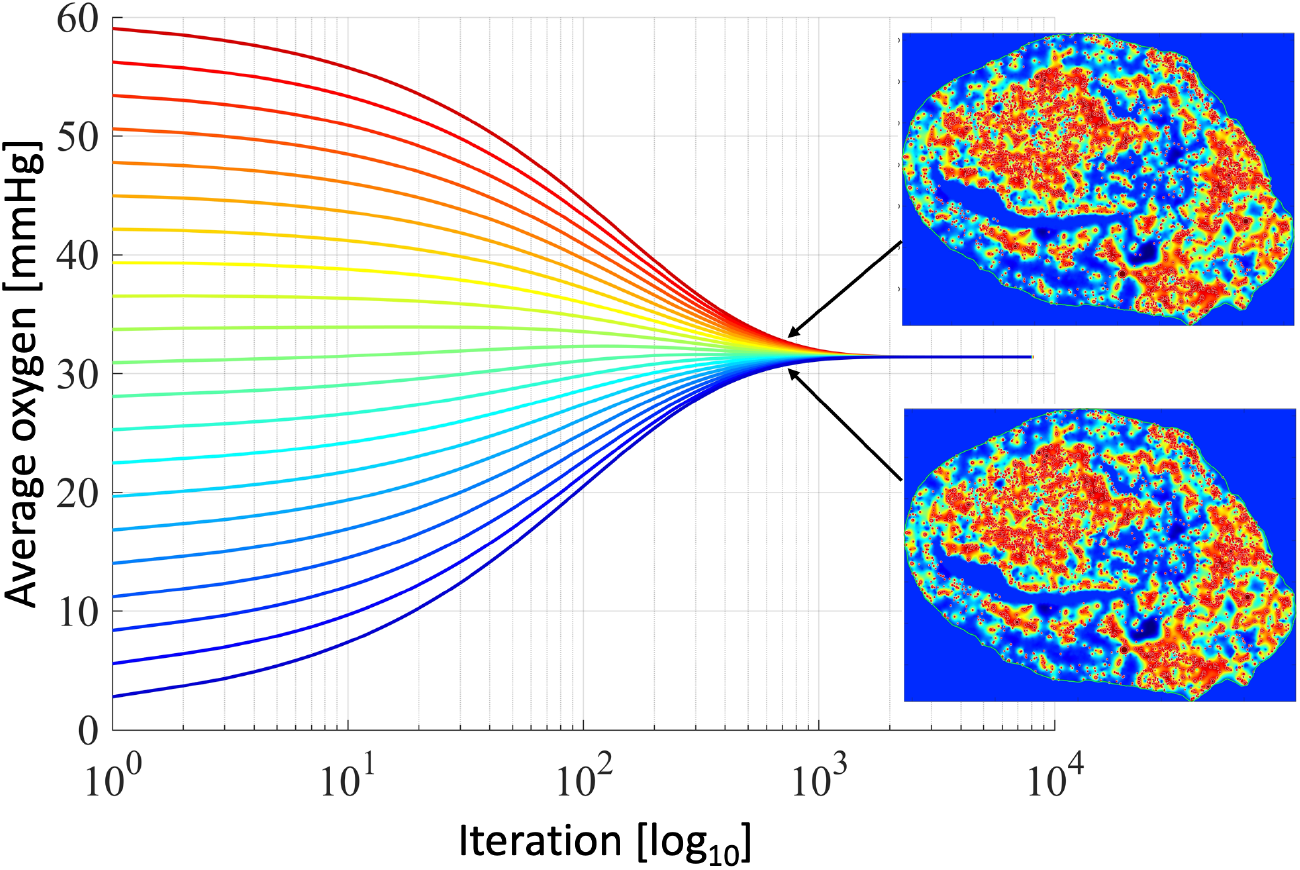
Independence of the final oxygen stabilization level on the initial oxygen amount in the tissue. Temporal evolution of the average oxygen levels in the tissue (y-axis) over iterations (x-axis, logarithmic scale) for 21 simulations initialized with 0 to 60 mmHg of oxygen, with increments of 3 mmHg. All graphs stabilized at 31.403 mmHg, with stabilization errors below 10^−5^. Each simulation is indicated by a different color corresponding to the initial tissue oxygenation level. The final stabilized oxygen maps are shown in the insets: a simulation initialized with 0 mmHg (bottom) and with 60 mmHg (top).

### 3.4. Sensitivity analysis of the domain boundary conditions

In the simulations discussed in sections 3.2 and 3.3, we assumed no loss or gain of oxygen along the computational domain boundaries and imposed the Neumann-type boundary conditions. These simulations resulted in the average oxygen level stabilizing at 31.403 mmHg after 7,981 iterations (Fig. 2). Here, we examined whether our model outcomes would change significantly if other types of boundary conditions were imposed. First, we tested the periodic boundary condition by replicating the oxygen values defined along each boundary column and row to the opposite side boundary. Simulations with periodic boundary conditions produced the same stabilized average oxygen level at 31.403 mmHg, and this also took 7,981 iterations. This stable oxygen distribution was not significantly different from that generated by the Neumann-type boundary conditions. The normalized L_2_ norm between the oxygen distributions was 2.619×10^−5^. This was calculated using only the grid points in the tissue and omitting the outer and cavity points, as they had the same oxygen level imposed (hypoxia). Next, we tested the Dirichlet boundary conditions with a constant hypoxia value defined along all the domain boundaries. The stabilized average oxygen level was again 31.403 mmHg, and the stabilization process required 7,981 iterations. The normalized L_2_ norm between solutions with the Dirichlet and Neumann boundary conditions was 2.619×10^−5^. The stable oxygen distributions produced by the Dirichlet and periodic boundary conditions were not significantly different from the stable oxygen map generated with the Neumann boundary conditions. Therefore, the final stable oxygen distribution was not sensitive to changing the domain boundary conditions because this condition is influenced by the outer points boundary condition discussed in 3.5.

### 3.5. Sensitivity analysis of the tissue boundary conditions

The histology image of the mouse bladder that we used as the base of our simulations did not occupy the whole rectangular computational domain. Therefore, we did not calculate oxygen diffusion in the grid points outside the tissue (outer points shown in magenta in Fig.4A). The boundary condition imposed on outer points affects the domain boundary conditions discussed in section 3.4. We imposed a constant hypoxia value in all these points as a boundary condition. Moreover, the bladder tissue contained some inner cavities (shown in blue in Fig. 4A). We also imposed hypoxia values in those grid points. These boundary conditions ensured that the inner cavities and outer points had constant hypoxia values during the simulation. We tested whether changing the boundary condition values from hypoxia (10 mmHg) to 0 mmHg would affect the final oxygen distribution within the tissue. Fig. 4B shows the stabilized average oxygen levels in the tissue: 31.403 mmHg for the hypoxia boundary condition and 28.536 mmHg for the zero-boundary condition. This gave a 2.867 mmHg difference between the averages. The normalized L_2_ norm between the final stabilized oxygen distributions was 0.0045. The L_2_ norm calculations used only the tissue grid points and were normalized by the number of tissue grid points. Therefore, the model output was sensitive to changing the oxygen level in the inner cavities and outer points. The stable oxygen maps for hypoxia and zero oxygen boundary conditions are shown in Figs. 4C and 4D, respectively.

**Fig. 4.**
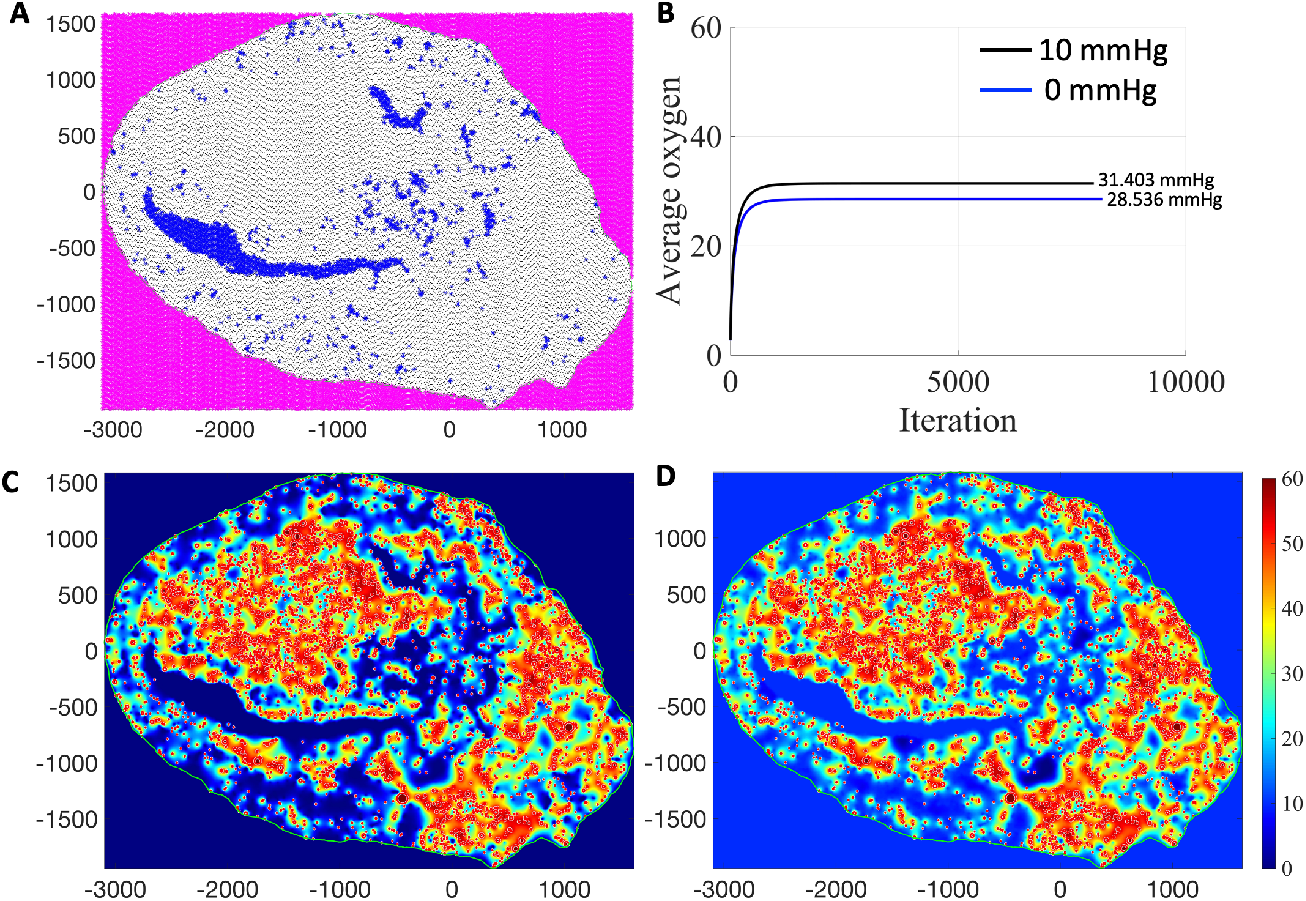
The dependence of the final stabilized oxygen distribution on the oxygen level imposed outside the tissue and in the inner cavities. **A**. Annotated grid points outside the tissue (magenta) and in the inner cavities (blue). The remaining area is composed of tissue grid points. **B**. Temporal evolution of the average oxygen levels with 0 mmHg (blue) and 10 mmHg (black) boundary conditions. **C**. The stabilized oxygenation map for 0 mmHg boundary condition. **D**. The stabilized oxygenation map for 10 mmHg boundary condition.

### 3.6. Sensitivity analysis of the vascular influx

In this section, we test the sensitivity of changing the influx parameter. The partial pressure of oxygen (pO_2_ in mmHg) in blood vessels varies depending on the vasculature type and size. For example, the pO_2_ in capillaries is 30 mmHg and in arteries is 80 mmHg [18, 23]. In this work, the base pO_2_ level in each vessel was set to 60 mmHg following [18, 23]. Therefore, we tested how robust the model outputs were to local perturbations when the influx was varied in a range of 60 +/-10 mmHg with a 1 mmHg progression. We investigated 20 parameter values and compared their model outputs with those generated by the baseline value of 60 mmHg.

The first model output analyzed was the cellular oxygen levels. We used *Â*-measure to assess the extent and significance of the changes in this model output. Fig. 5A shows the *Â*-measure values (y-axis) calculated by comparing the cellular oxygen levels produced by the perturbed values vs. those produced by the baseline value. The plot shows that the influx within 4 mmHg from 60 mmHg yielded model outputs that were not statistically different from the baseline value—they are within the green lines in Fig. 5A, as indicated by ‘*’, and thus they are within the minimal statistical significance as determined by the A-measure method.

**Fig. 5.**
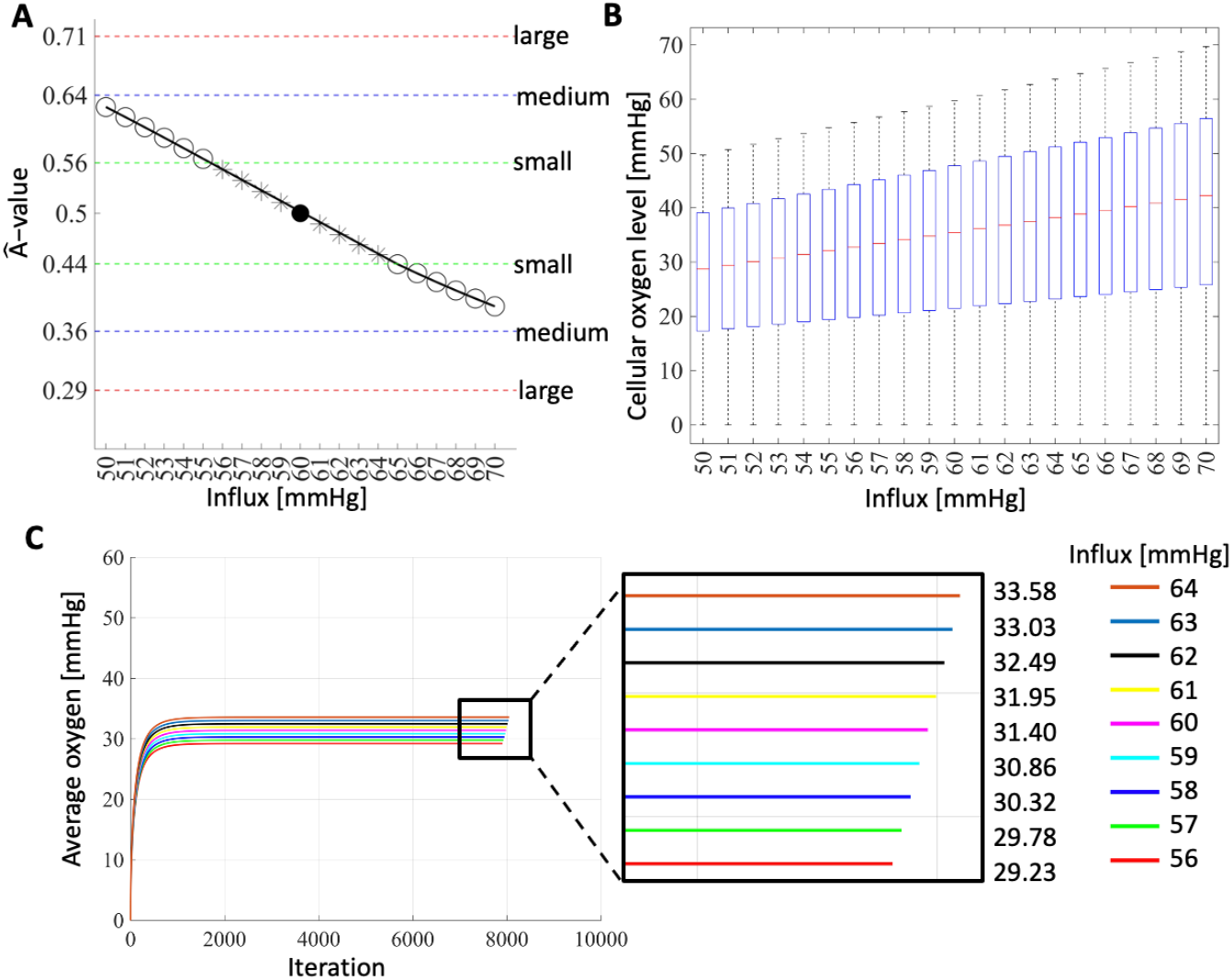
The model outputs are independent of the vascular influx value for small perturbations. **A**. The *Â*-values (y-axis) calculated by comparing the cellular oxygen levels produced by perturbed and the baseline influx values (x-axis). The symbols represent: filled circle (baseline value), stars (perturbed values plotted for panel C), and open circles (other perturbed values). **B**. Boxplots of the cellular oxygen levels (y-axis) for all perturbed and the baseline influx values (x-axis) from A. **C**. Temporal evolution of the average tissue oxygen levels (y-axis) over iterations (x-axis). The inset shows a magnified portion with the corresponding average oxygen values.

Thus, the cellular oxygen levels are less sensitive to +/-4 mmHg perturbations of the vascular influx. However, influx values beyond 4 mmHg perturbations produced model outputs that differed significantly from the baseline value. These sensitive parameter values are outside the green lines in Fig. 5A. Fig. 5B presents the five-point boxplots of the cellular oxygen levels for the baseline value, and all considered perturbed values from Fig. 5A. The boxplots show that the median cellular oxygen levels increased as the influx values increased. As expected, the minimum cellular oxygen levels were zero because some cells do not have oxygen. In contrast, the maximum cellular oxygen levels corresponded to the influx value used in the simulation.

The second model output analyzed was the average oxygen levels in the tissue in each iteration. Fig. 5C shows this output plotted for the perturbed values indicated by ‘*’ in Fig. 5A and the baseline value. The average oxygen distribution stabilized at levels from 29.234 mmHg to 33.577 mmHg for the influx values between 56 mmHg and 64 mmHg, with about 0.543 mmHg difference for each 1 mmHg increment of the influx value. These close stabilizations support the conclusion drawn from Fig 5A. The number of iterations required to reach the stabilized oxygen distribution varied depending on the influx value. It was fastest for 56 mmHg and increased with each additional 1 mmHg increment until 64 mmHg.

The third model output analyzed was the final stabilized oxygen distribution in the tissue. The L_2_norms between this output generated by the baseline value and those generated for each perturbed value were calculated. The minimum and maximum normalized L_2_ norm values were 0.0008 and 0.0083, respectively. This shows that as the parameter was perturbed, the final stable oxygen distribution did not change significantly compared to that generated by the baseline parameter.

### 3.7 Sensitivity analysis of the maximum cellular uptake rate *V*_*m*_

We used the Michaelis–Menten kinetics to model the cellular uptake of oxygen. However, the consumption rate 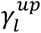 was defined by Eqs. (13) or (15), depending on the cell size. The Michaelis–Menten parameter values for *V*_*m*_, the maximum oxygen consumption rate, and *K*_*m*_, the oxygen concentration at half consumption rate, were initially estimated based on [24]. The *V*_*m*_ value was further optimized for this tissue, as described below. As chronic hypoxia in tumors typically occurs at distances between 100 and 180 μ*m* from the vasculature [2], the cells in our tissue whose minimum Euclidean distance from the closest vessel was greater than 100 μ*m* were designated as hypoxic candidates (shown in green in Fig. 1C). For the tissue under consideration, they constituted 0.52 % of all cells (266 out of 51266). The *patternsearch* routine in MATLAB®, a direct search algorithms, was used to find the baseline *V*_*m*_ value. Our objective function was set up to minimize the false negatives (the cells that are hypoxic candidates and not simulated as hypoxic). Initially, *V*_*m*_=6.25×10^−17^ mole/s per cell [24] (=9.548 σg/μm^3^*s after rescaling). Simulations with this parameter value yielded an accuracy of 0.86, a misclassification rate of 0.14, a sensitivity of 1, and a specificity of 0.86. The optimized baseline value was 3.125×10^−17^ mole/s per cell (=4.774 σg/μm^3^*s), resulting in an accuracy of 0.96, a misclassification rate of 0.04, a sensitivity of 0.89, and a specificity of 0.96. We then tested the local sensitivity around this optimized baseline value using 20 perturbed values. First, the extent and significance of the changes in the cellular oxygen levels are summarized in the *Â*-measure plot in Fig. 6A. The plot shows that changing *V*_*m*_ between 1.64×10^−17^ and 4.9×10^−17^ mole/s per cell yielded model outputs whose differences compared to the baseline value outputs were small. The statistical significance of the differences was minimal, meaning that the cellular oxygen levels were less sensitive to these perturbations (indicated by * within the green lines in Fig. 6A). These are the robust values of *V*_*m*_. The cellular oxygen levels were more sensitive to perturbed *V*_*m*_ values outside this range. Second, the boxplots of the oxygen levels sensed by the cells are shown in Fig. 6B. For smaller *V*_*m*_ and thus smaller consumption, the median cellular oxygen level was increased in/around the cells because there was more oxygen left in the microenvironment. In contrast, larger *V*_*m*_ implies more consumption and less oxygen left in the microenvironment. Third, Fig. 6C supports this observation by showing the inverse relationship between *V*_*m*_ and the average oxygen level in the tissue. As expected, the stabilized average oxygen value increased as the *V*_*m*_ decreased because cells consumed less oxygen, and more oxygen remained in the interstitial space of the tissue. Finally, the normalized L_2_ norm values were calculated for the final stabilized oxygen distributions produced by the perturbed values compared to those produced by the baseline value. The minimum and maximum normalized L_2_ norm values were 0.0003 and 0.0062, respectively.

**Fig. 6.**
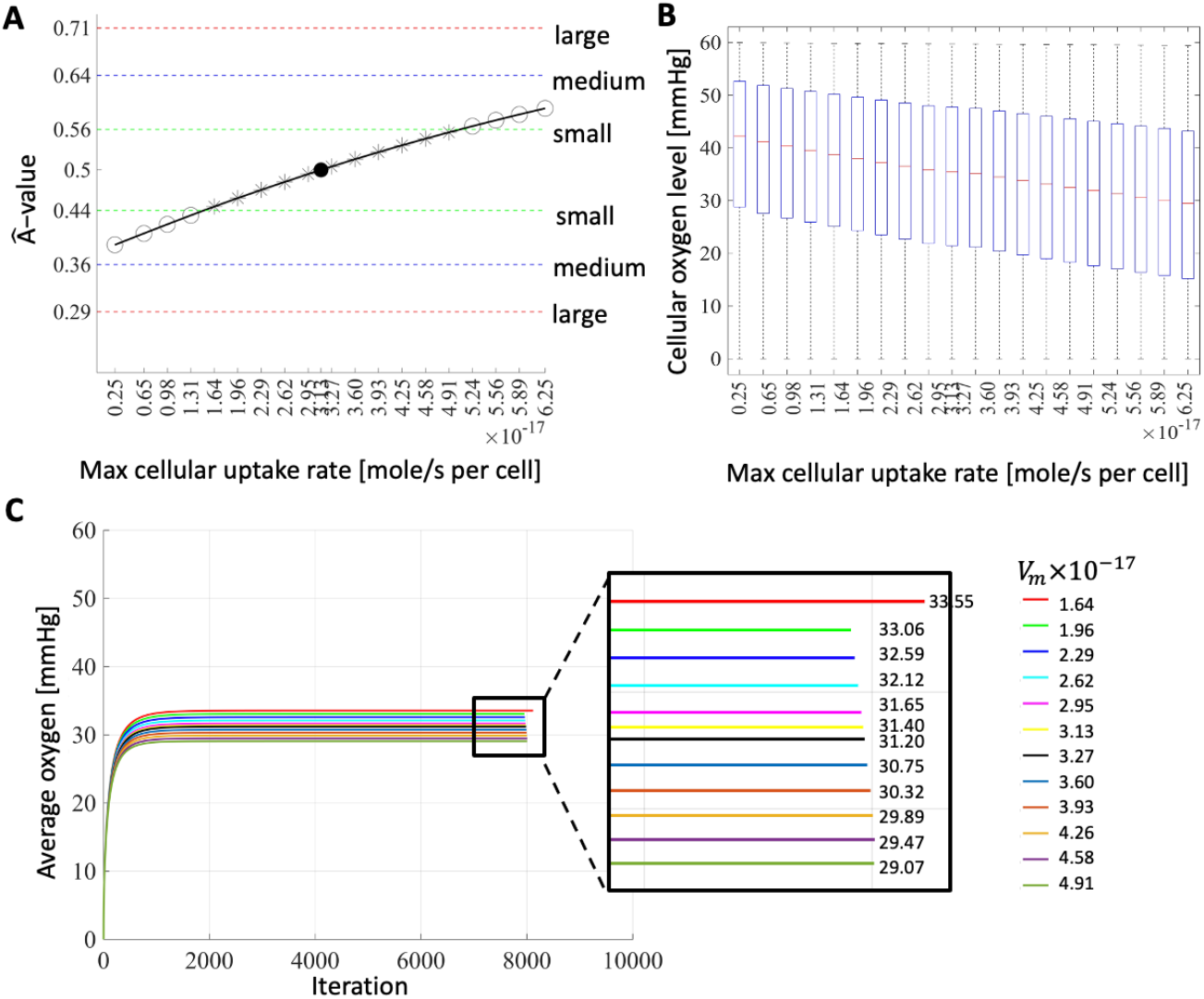
Sensitivity analysis to *V*_*m*_ parameter. **A**. The *Â*values (y-axis) calculated by comparing the cellular oxygen levels produced by perturbed and the baseline *V*_*m*_ values (x-axis). The symbols represent: filled circle (baseline value), stars (perturbed values plotted for panel C), and open circles (other perturbed values). **B**. Boxplots of the cellular oxygen levels (y-axis) for the perturbed and the baseline *V*_*m*_ values (x-axis) from A. **C**. Temporal evolution of the average oxygen in the tissue (y-axis) over iterations (x-axis). The inset shows a magnified portion with the corresponding average oxygen values.

First, the extent and significance of the changes in the cellular oxygen levels are summarized in the *Â*-measure plot in Fig. 6A. The plot shows that changing *V*_*m*_ between 1.64×10^−17^ and 4.9×10^−17^ mole/s per cell yielded model outputs whose differences compared to the baseline value outputs were small. The statistical significance of the differences was minimal, meaning that the cellular oxygen levels were less sensitive to these perturbations (indicated by * within the green lines in Fig. 6A). These are the robust values of *V*_*m*_. The cellular oxygen levels were more sensitive to perturbed *V*_*m*_ values outside this range. Second, the boxplots of the oxygen levels sensed by the cells are shown in Fig. 6B. For smaller *V*_*m*_ and thus smaller consumption, the median cellular oxygen level was increased in/around the cells because there was more oxygen left in the microenvironment. In contrast, larger *V*_*m*_ implies more consumption and less oxygen left in the microenvironment. Third, Fig. 6C supports this observation by showing the inverse relationship between *V*_*m*_ and the average oxygen level in the tissue. As expected, the stabilized average oxygen value increased as the *V*_*m*_ decreased because cells consumed less oxygen, and more oxygen remained in the interstitial space of the tissue. Finally, the normalized L_2_norm values were calculated for the final stabilized oxygen distributions produced by the perturbed values compared to those produced by the baseline value. The minimum and maximum normalized L_2_ norm values were 0.0003 and 0.0062, respectively.

### 3.8. Sensitivity analysis of the Michaelis constant *K*_*m*_

The value of the Michaelis constant *K*_*m*_ was estimated in [24] to be *K*_*m*_= 2.1×10^−3^ mM of O_2_ (=1.344 *σg*/μ*m*^3^ after scaling). *K*_*m*_ is inversely related to the affinity for oxygen binding; thus, for high *K*_*m*_ values, the cells require greater oxygen concentrations to reach the maximal consumption rate *V*_*m*_. In contrast, low values of *K*_*m*_ show a high affinity for oxygen binding and require lower oxygen concentrations to reach *V*_*m*_. We conducted a value sensitivity analysis for *K*_*m*_ by dividing the baseline value (by two, four, and ten) to obtain small values and multiplying the baseline value (by the same numbers and more) to obtain large values. Therefore, we tested 13 parameter values.

First, the magnitude and significance of the changes in cellular oxygen level produced by the perturbed vs. baseline value are summarized in Fig. 7A *Â*-measure plot. The plot shows that none of the perturbed values tested resulted in significantly different model outputs compared to the baseline value output, meaning, the cellular oxygen levels are insensitive to *K*_*m*_ perturbations. A large *K*_*m*_= 73.5×10^−3^ value was required to cross the green line. Second, the boxplots of the cellular oxygen levels in Fig. 7B show that for smaller values of *K*_*m*_, the median cellular oxygen levels were the same. However, the median values increased starting at *K*_*m*_= 21×10^−3^. The minimum and maximum cellular oxygen values were the same for all values. This is because some cells will not uptake oxygen while others will uptake oxygen up to the imposed vascular influx value.

**Fig. 7.**
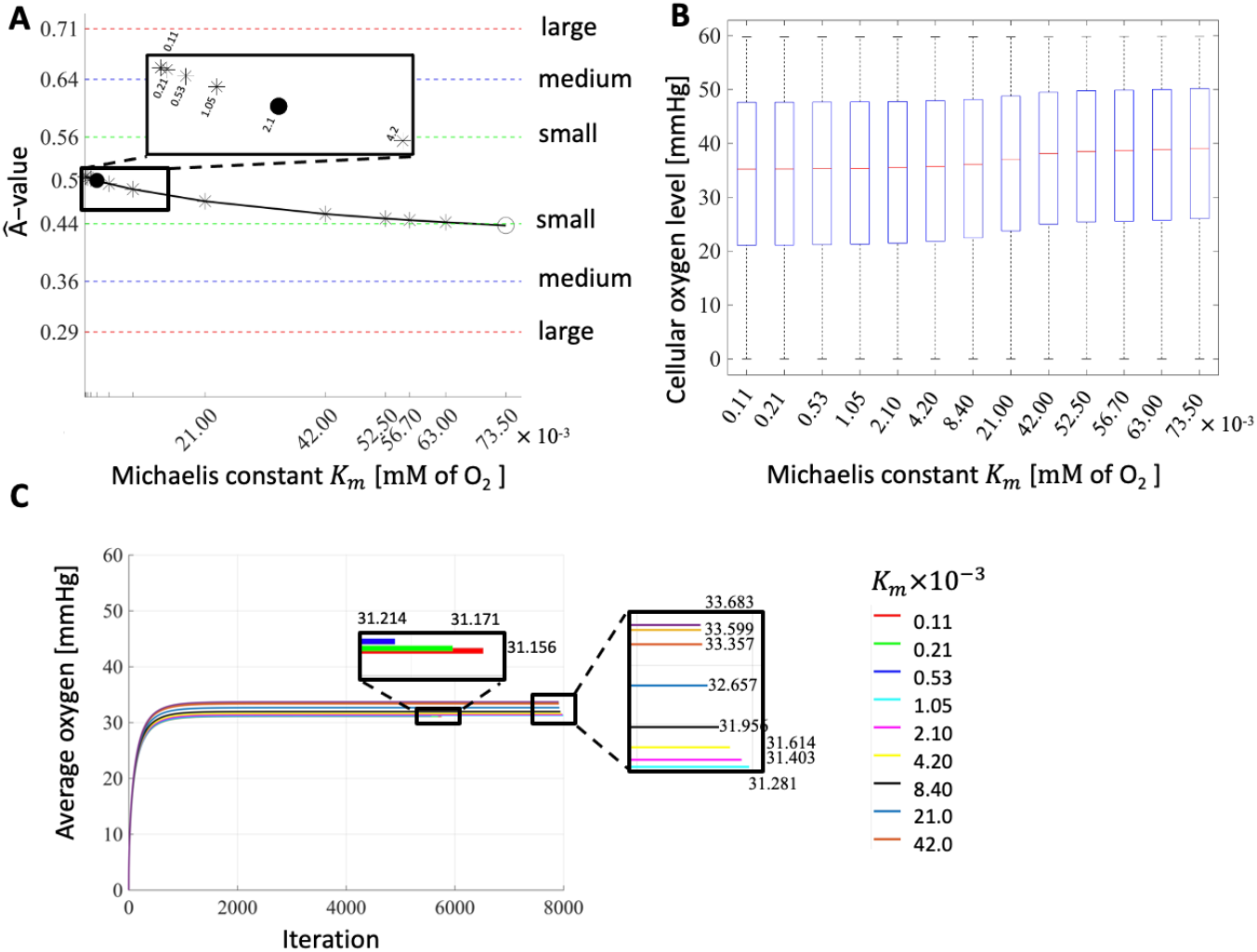
Sensitivity analysis to *K*_*m*_ parameter. **A**. The *Â*-values (y-axis) were calculated by comparing the cellular oxygen levels produced by perturbed and baseline *K*_*m*_ values (x-axis). The symbols represent: filled circle (baseline value), stars (perturbed values plotted for panel C), and open circles (other perturbed values). It takes *K*_*m*_ to be a large as 73.5 ×10^−3^ mM of O_2_ to cross the green line). **B**. Boxplots of the cellular oxygen levels (y-axis) for the perturbed and the baseline *K*_*m*_ values (x-axis) from A. **C**. Temporal evolution of the average oxygen in the tissue (y-axis) over iterations (x-axis) for simulations with *K*_*m*_ between 0.11 ×10^−3^ and 42 ×10^−3^ mM of O_2_. The two insets show zoomed-in portions with the corresponding average oxygen values.

Third, the average oxygen in the tissue for each value was calculated and plotted in Fig 7C. This graph shows that the average oxygen levels increased as the Michaelis constant *K*_*m*_ increased. Larger *K*_*m*_ values imply low affinity for oxygen binding, resulting in more oxygen in the tissue and a higher average oxygen value. Conversely, lower *K*_*m*_ values result in a lower average oxygen level. However, the averages were close to each other. In addition, the normalized L_2_ norm values between the final stabilized oxygen gradients for perturbed values compared to the baseline value were 0.0002 and 0.0039 for the minimum and maximum values, respectively. These small values indicate that the stabilized oxygen gradients were not significantly different from the map generated by the baseline value. Therefore, the model outputs were less sensitive to both small and large perturbations of *K*_*m*_.

## 4. DISCUSSION

In this paper, we presented a mathematical model for reconstructing the oxygenation landscape of a digitized tissue obtained from *in vivo* murine experiments. Similar mathematical models were used previously by our group [3, 10, 11]. However, in the current study, we performed computational simulations using an actual image of a mouse bladder tissue and tumors. Most parameters in our study were based on published experimental measurements (summarized in Table 1). However, like all experimental measurements, these parameters were associated with some variability and uncertainty. Therefore, we used sensitivity analyses to examine how changes in model parameters would affect the outcomes of the mathematical model. These analyses helped to determine the robust parameters and their changeable ranges vs. those that are sensitive and should be fixed.

Our sensitivity analysis of the initial oxygen level in the tissue shows that this initial condition is robust. We also observed that the simulations that started with lower oxygen levels stabilized more quickly than those with higher initial oxygen levels. These results corroborate those of Kingsley et al. [3], who found, that the final stabilized oxygen distribution is independent of the oxygen level chosen to initiate the oxygenation process, but some simulations stabilized faster than others. However, our work differs in several key ways from [3]. Specifically, the vasculature in [3] comprised vessels of the same size and identical oxygen influx levels. Similarly, all cells of the same kind (tumor or stromal) had identical sizes. In contrast, the structural tissue elements in our model were of variable sizes, with non-uniform vascular influx and cellular consumption modes based on cell size. Furthermore, the tissue in [3] was smaller in size, had a square shape, and lacked some of our tissues’ features, such as inner cavities and an irregular tissue shape that required the definition of outer points. These features make our model more realistic and bring it closer to translation from preclinical to clinical work and validation with patients’ histological-based tissues. Our next sensitivity analysis showed that changing the type of tissue boundary conditions did not affect the final model output. Using the periodic or Dirichlet instead of Neumann boundary conditions on the domain boundaries did not significantly change the final stabilized oxygen distribution. However, the model was sensitive to changing the imposed hypoxia values in the inner cavity’s points. The hypoxia boundary condition is biologically feasible given that these cavities (representing the hollow spaces of the bladder) may be exposed to some oxygen level, albeit a low-level [25]. Therefore, we recommend imposing the hypoxia boundary condition. The high sensitivity to the changes in cavity values from hypoxia to zero indicates that this parameter may need to be measured experimentally to improve the model’s accuracy. The natural levels of oxygen partial pressure may be assessed in such cavity spaces using direct oxygen imaging with solid probes, as proposed in [26].

The sensitivity analysis of the oxygen vascular influx level showed that this parameter is robust only for the oxygen pressure gradients within 4 mmHg from the baseline value, which was set to 60 mmHg [18, 23]. The parameter values from 56 to 64 mmHg produced results with minimal statistically significant differences compared to the baseline value. However, the influx parameter is sensitive to changes beyond 4 mmHg, and thus it should be measured experimentally for a given tumor, as tumor vascularization may differ for different cancers. Our study also suggests that of the two Michaelis–Menten parameters, the model is more sensitive to large perturbations of the maximum consumption rate *V*_*m*_. Only small perturbations of *V*_*m*_ have a negligible influence on the final stabilized average oxygen levels and the resultant cellular oxygen levels. Conversely, no perturbations of *K*_*m*_, the oxygen concentration at half *V*_*m*_, produced significantly different results compared to the baseline value. Our results are consistent with other sensitivity analyses by Fontes et al. [27] and Hetrick et al. [28], who showed that their models were also sensitive to *V*_*m*_ and insensitive to *K*_*m*_.

The tissue oxygenation in our model was recreated based on 2-dimensional (2D) histology images, whereas truly 3-dimensional (3D) recreation would be useful. The 2D tissue slices are important as they provide information on cellularity, vascularity, and spatial heterogeneity and lead to identifying subregions with different oxygenation patterns. Nevertheless, 3D models offer more realistic and detailed geometries, easier visualization of tumor volumetric data, and may also provide more physiologically accurate information. Our model can be translated to the 3D space by using spheres (of different radii) to represent the cells, as was done in [29, 30], and branched tube segments that can represent vasculature or irregular geometry, as was done in [31-33]. The equations can be extended to 3D by including the z plane. For example, the spatial locations of 3D grid points would be ***x*** = (*x, y, z*) as was done in [34]. However, visualizing the diffused oxygen gradients may be difficult in 3D. Furthermore, these 3D models may be computationally expensive.

## CONCLUSIONS

In this study, we analyzed the robustness of the physical and computational parameters in our hybrid ABM. Based on a local sensitivity analysis, we identified that the most robust parameters are: the Michaelis constant *K*_*m*_, the computational domain boundary conditions, and the initial oxygen conditions. Conversely, the maximum consumption rate *V*_*m*_, the vascular influx, and the tissue boundary condition imposed on the outer points and the inner cavities are sensitive parameters.

## ACKNOWLEDGMENTS

This work was supported in part by the CA259387 grant from the National Institutes of Health and in part by the Moffitt Tissue Core and Moffitt Analytic Microscopy Core facilities at the Moffitt Cancer Center under the Cancer Center Support Grant P30-CA076292 from the National Cancer Institute.

